# A functional single nucleotide polymorphism upstream of the collagen type III gene may contribute to catastrophic fracture risk in Thoroughbred horses

**DOI:** 10.1101/2023.06.16.545331

**Authors:** Esther Palomino Lago, Arabella Baird, Sarah C. Blott, Rhona E. McPhail, Amy C. Ross, Sian A. Durward-Akhurst, Deborah J. Guest

## Abstract

Fractures caused by bone overloading are a leading cause of euthanasia in Thoroughbred racehorses. The risk of fatal fracture has been shown to be influenced by both environmental and genetic factors but, to date, no specific genetic mechanisms underpinning fracture have been identified. The aim of this study was to utilise a genome-wide polygenic risk score to establish an *in vitro* cell system to study bone gene regulation in horses at high and low genetic risk of fracture. Candidate gene expression analysis revealed differential expression of *COL3A1* and *STAT1* genes in osteoblasts derived from high and low risk horses. Whole genome sequencing of fracture case and control horses revealed a single nucleotide polymorphism (SNP) upstream of *COL3A1* that was confirmed in a larger cohort to be significantly associated with fracture. Bioinformatics tools predicted that this SNP may impact the binding of the transcription factor SOX11. Gene modulation demonstrated SOX11 is upstream of *COL3A1* and the region binds to nuclear proteins. Furthermore, luciferase assays demonstrated that the region containing the SNP has promoter activity. However, the specific effect of the SNP depends on the broader genetic background of the cells and suggest other factors may also be involved in regulating *COL3A1* expression. In conclusion, this approach provides a powerful means to identify novel DNA variants and understand their mechanism of action to enable the development of new ways to identify and treat horses at high risk of a catastrophic fracture.

## Introduction

Bone fractures caused by overloading rather than direct trauma can occur in Thoroughbred racehorses and, due to the complexities involved in treating fractures in a large animal that must remain weight bearing, many of these can be catastrophic and result in euthanasia. Bone fractures are the leading cause of deaths occurring on the racecourse (McKee 1995) and therefore have a significant welfare and economic impact on the racing industry. Many environmental risk factors have been identified for catastrophic fracture (Parkin *et al*. 2004; Verheyen *et al*. 2006; Anthenill *et al*. 2007; Kristoffersen *et al*. 2010; Georgopoulos & Parkin 2017) but there is also a genetic contribution to risk. The heritability of distal limb fracture has been found to range from 0.21-0.37 (Welsh *et al*. 2014) and multiple genomic locations have been identified that may contribute to fracture risk, including a 5 Mb region on equine chromosome 18 (ECA18) (Blott *et al*. 2014; Blott & Swinburne 2015). A candidate SNP study using SNPs within this region also confirmed an association with fracture (Tozaki *et al*. 2020). Similarly both genetic (Korvala *et al*. 2010; Varley *et al*. 2016; Zhao *et al*. 2016; Bulathsinhala *et al*. 2017) and environmental (Warden *et al*. 2006; Warden *et al*. 2007) risk factors have been identified for stress fractures in humans. However, fracture risk is a complex trait resulting from the effects of multiple genes and no specific functional mechanisms underpinning fracture have been defined in either species. To enable better prevention of catastrophic fractures, we need to develop improved understanding of the biological processes affected by genetic factors.

The 5 Mb region on ECA 18 that is associated with catastrophic fracture risk in horses (Blott *et al*. 2014) contains eleven genes that are involved in fracture or bone formation: *ZNF804A* (Long *et al*. 2012; Blott *et al*. 2014), *ITGAV* (Logan *et al*. 2018), *CALCRL* (Wang *et al*. 2010), GULP1 (Park *et al*. 2016), *COL3A1* (Volk *et al*. 2014), *COL5A2* (Hong *et al*. 2010), *SLC40A1* (Pereira *et al*. 2020), *MSTN* (Bialek *et al*. 2014; Wu *et al*. 2018), *C2orf88* (Burger *et al*. 2017), *STAT1* (Kim *et al*. 2003; Xiao *et al*. 2004; Tajima *et al*. 2010), *GLS* (Yu *et al*. 2019). Further investigation of these genes in equine fracture is therefore warranted.

However, this may be difficult to do in horses *in vivo*, due to the large variation in exposure to environmental factors of fracture risk (e.g. training and racing histories) and the fact that bone tissue can only be isolated post-mortem. Small animal models are very valuable for defining gene functions (Groza *et al*. 2022), but are unlikely to be useful in studying specific DNA variants as causal variants are often poorly conserved between species (Flint & Mackay 2009). Cell models have been widely used to study human diseases (Li & Zhou 2010). Often these are simple inherited diseases but for complex genetic diseases cells from affected patients have been compared to cells from unaffected donors (Park *et al*. 2008; Barral & Kurian 2016; Chamberlain 2016). However, more recently these models have been refined by the selection of cells based on their polygenic risk score (Dobrindt *et al*. 2021; Coleman 2022). In contrast to humans, the use of cell models to study inherited diseases in horses has not been widely reported.

The aim of this study was to apply genome-wide polygenic risk scores (Genomic Best Linear Unbiased Prediction, GBLUP) (Choi *et al*. 2020; Chung 2021) with *in vitro* disease modelling in horses to investigate the regulation of candidate genes in horses at genetically high and low risk of fracture.

## Materials and Methods

An overview of the experimental approach is shown in Figure 1.

**Figure 1.**
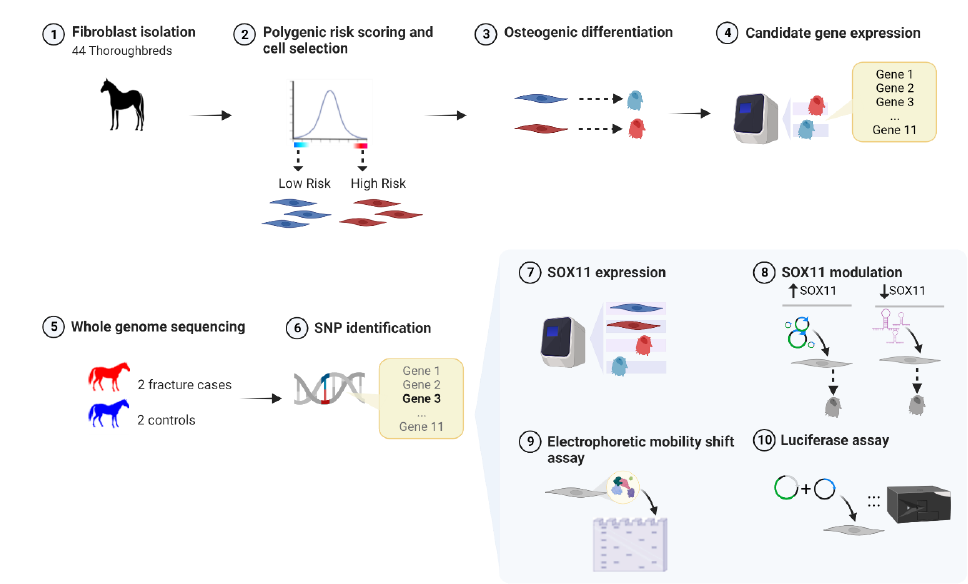
An overview of the experimental approach used demonstrating the order of the experiments. Created with BioRender.com

### Skin fibroblast cell isolation and culture

Equine skin fibroblasts were isolated post-mortem from 44 Thoroughbred horses (of unknown fracture status) and three Welsh Mountain ponies. All animals had been euthanised for reasons unrelated to this study and with the consent of the Animal Health Trust Ethical Review Committee (AHT_02_2012) and Royal Veterinary College Clinical Research Ethical Review Board (URN 2021 2035-2). Briefly, small dermal tissue pieces were collected into standard culture media (Dulbecco’s modified Eagle’s medium (DMEM) containing 10% fetal bovine serum, 2 mM L-glutamine, 100 U/ml penicillin, 100 µg/ml streptomycin (all ThermoFisher, Loughborough, UK) plus 2.5 µg/ml amphotericin B and stored at 4°C for up to 24 h. Tissue samples were then dissected into small pieces and digested in 1 mg/ml collagenase from *C. histolyticum* (Sigma-Aldrich, Dorset, UK) in standard culture media plus 2.5 µg/ml amphotericin B overnight at 37°C, 5% CO_2_. Dissociated cells were washed and cultured in standard culture media (without amphotericin B) and passaged using 0.25% trysin-EDTA (Sigma-Aldrich) every 3-5 days.

### Polygenic risk scoring of Thoroughbred skin fibroblasts

Genome wide Complex Traits Analysis (GCTA) (Yang *et al*. 2011) was used on Equine SNP50 genotyping data from 269 cases and 253 controls from the previous study (Blott *et al*. 2014) (“the training data set”) to derive BLUP (Best Linear Unbiased Prediction) allele effects for all individual SNPs across the genome (genomic BLUP, GBLUP). This is equivalent to deriving the heritability explained by each individual SNP, a pre-requisite for polygenic risk score analysis (Choi *et al*. 2020). DNA was extracted from skin fibroblasts using the QIAamp DNA mini kit (Qiagen, Manchester, UK) and genotyped on the Geneseek Genomic Profiler Equine Plus chip (based on Illumina’s Equine SNP70 platform) (performed by Neogen, Ayr, UK), which includes over 70,000 evenly distributed SNPs including those on the Illumina Equine SNP50 chip. Polygenic risk scores for the skin fibroblast samples were calculated by applying the PLINK (http://pngu.mgh.harvard.edu/purcell/plink/ (Purcell *et al*. 2007)) score function to the fibroblast genotypes, in conjunction with the estimated SNP BLUP effects obtained from the training data set, applying the GBLUP protocol as described by Chung (Chung 2021). Additionally, we carried out genotyping and scoring of 382 DNA samples which had been collected from blood samples taken from Thoroughbreds of unknown fracture status. These Thoroughbreds therefore represent the general population to enable us to select skin samples that represented the ends of the risk spectrum. Skin fibroblasts from only male horses were selected for use in this study.

### Osteoblast differentiation

To promote osteoblast-like cell differentiation of the primary equine fibroblasts cells at approximately 70% confluency were cultured in osteogenic media (standard culture media plus 10 nM dexamethasone, 28 µM ascorbic acid and either 2 mM or 10 mM β-glycerophosphate, (all Sigma-Aldrich) for 21 days, with the media replaced every 3-4 days.

### Histological staining

To determine matrix mineralisation following osteoblast-like cell differentiation, cells were stained with von Kossa (Abcam, Cambridgeshire, UK) according to the manufacturer’s instructions. Calcium deposition was detected by incubating the cells with 2% Alizarin red S pH4 4.2 for 5 min. Hydroxyapatite deposition was detected using the OsteoImage bone mineralisation assay (Lonza, Slough, UK).

### RNA extraction, cDNA synthesis and quantitative PCR

RNA was extracted from skin cells (pre and post osteogenic differentiation and following SOX11 overexpression and knockdown) using Tri-reagent (Sigma-Aldrich) and purified using an RNeasy mini kit (Qiagen). Genomic DNA contamination was removed using Invitrogen™ DNA-free™ DNA Removal Kit (ThermoFisher). 1 µg of RNA was used to prepare cDNA using the Sensifast cDNA synthesis kit (Bioline, London, UK). 2 µl aliquots of cDNA (corresponding to 20 ng of cDNA) were used in qPCR using SYBR green containing supermix (Bioline) on the Biorad CFX96 thermal cycler (Biorad, Hertfordshire, UK) in duplicate. Primers to equine specific genes (note SOX11 primers detect both the human and equine genes) were designed using NCBI primer blast (https://www.ncbi.nlm.nih.gov/tools/primer-blast/) and mfold (http://www.unafold.org/) programs to produce amplicons of 50-150 bp, a melting temperature ™ of 58-62°C and devoid of secondary structures at Tm 60°C. Primer sequences are in Table S1. qPCR cycling parameters were 95°C for 10 min, followed by 45 cycles of 95°C for 15 s, 60°C for 15s and 72°C for 15s. Gene expression was measured relative to the 18s rRNA housekeeping gene using the 1/2^ΔΔCT^(Livak & Schmittgen 2001). The 18s rRNA was selected as a housekeeping gene based on its stability across the samples and experimental conditions being tested. Stability of 18s rRNA, GAPDH and ACTB housekeeping genes were evaluated across the experiments using RefFinder (Xie *et al*. 2012) (https://www.heartcure.com.au/reffinder, Supplementary Figure 1).

### Immunocytochemistry

Skin fibroblasts pre and post osteogenic differentiation were cultured on gelatin coated coverslips (Sigma-Aldrich), fixed in 3% paraformaldehyde for 20 min and permeabilised for 1 h with 0.1% triton-X-100 prior to blocking in 2.5% normal horse serum (Vector Laboratories, Peterborough, UK) for 20 min. Primary antibody incubations were performed at 4°C overnight prior to incubation with the secondary antibody. All antibodies were used at optimised concentrations in blocking solution (Table S2). Coverslips were mounted onto glass sides using Vectashield Hardset mounting media with DAPI (Vector Laboratories) and imaged using a Nikon Eclipse Ti2 series microscope (Nikon, Surrey, UK). For RUNX2 and STAT1 staining, the intensity of nuclear staining was quantified using ImageJ (Schneider *et al*. 2012).

### Whole Genome Sequencing

Whole genome sequencing was performed on two Thoroughbred fracture cases and two controls. The DNA samples were isolated as part of a previous study (Blott *et al*. 2014). The fracture cases sustained distal limb fractures (one horse had a sesamoid fracture, one horse had a lateral condylar fracture) that required euthanasia while racing on a UK racecourse. The fracture controls were uninjured horses over four years of age with no history of fracture and had been racing during the same time period as the cases. Whole genome sequencing was performed by Source Bioscience (Cambridge, UK) on 100 bp (base pair) PE (paired end) reads using an Illumina HiSeq 2000 to generate 30-36x coverage per sample. Reads were aligned to the reference genome (Equicab 3.0) and variant calls made using GATK (HaplotypeCaller) (McKenna *et al*. 2010). Base quality score recalibration, and indel realignment and duplicate removal were carried out and SNP discovery performed according to GATK Best Practices (Van der Auwera *et al*. 2013). Genomic Variant Call Format (VCF) files were combined into a single VCF file and Variant Effect Predictor (VEP) (McLaren *et al*. 2016) used for cross-genome analysis. The effect of SNPs on transcription factor binding were predicted using TomTom (Gupta *et al*. 2007). The minor allele frequencies were calculated from publicly available datasets including: horses representing multiple breeds (http://gong_lab.hzau.edu.cn/Animal_SNPAtlas (Jagannathan *et al*. 2019)), 101 Japanese Thoroughbreds (Tozaki *et al*. 2021) and other individual breeds, including US/European Thoroughbreds (Durward-Akhurst *et al*. 2021). Genotypes of five SNP heterozygotes and five SNP homozygotes were confirmed using the Integrative Genome Viewer (Robinson *et al*. 2011). Histone modifications in the region containing the SNP were confirmed using ENCODE data (Luo *et al*. 2020) and FANNGmine (version 1.3, https://faangmine.rnet.missouri.edu/).

### Genotyping for the COL3A1 SNP

Allelic discrimination assays were performed on 86 fracture controls and 91 fracture cases, isolated as part of the previous study (Blott *et al*. 2014) (i.e. part of the discovery cohort used to develop the “training data set”). Reactions were carried out in 8 µl volumes consisting of 2 µl (100 ng) genomic DNA in 0.2 µl of 40X assay mix, 4 µl Luna universal qPCR Master Mix (New England Biolabs, Hitchin, UK) and 1.8 µl ultrapure water. Cycling parameters were 25°C for 3 seconds, 95°C for 3 minutes, followed by 35 cycles of 95°C for 3 seconds and 60°C for 10 seconds, with a final extension 25°C for 30 seconds. The primer and probe sequences were: 5′-TCTTGAGGAATTGGTCACAGGA −3′ (forward primer); 5′-AGGACTTGGGCATATCTAACCA −3′ (reverse primer); CAAATATT<u>T</u>GCCTGTCAGGTCCCACC (VIC, 4,7,2′-trichloro-7′-phenyl-6-carboxyfluorescein); and CAAATATT<u>G</u>GCCTGTCAGGTCCCACC (FAM, 6-carboxyfluorescein). Allelic discrimination was performed using an Applied Biosystems StepOnePlus™ Real-Time PCR system (ThermoFisher) and results were analysed using Applied Biosystems StepOne Software v2.3.

### Overexpression of SOX11

1×10^6^ skin fibroblasts (from one Thoroughbred and two Welsh Mountain ponies) were suspended in 400 µl of standard culture media lacking penicillin/ streptomycin and either 20 µg of pCMV6-AC-GFP human SOX11 (NM_003108; Origene, Maryland, US) or pCAG-eGFP (Liew *et al*. 2007) which was a gift from Peter Andrews (University of Sheffield, UK). Electroporation was performed in a 4 mm gap cuvette with one 35 ms pulse of 170 V using a BTX EM830 (BTX, San Diego, CA). 48h after electroporation, antibiotic resistance was used to select stably transfected cells using 1 mg /ml G418 or 3.5 µg/ml puromycin (both Sigma-Aldrich) respectively, for up to 14 days. Following selection, undifferentiated cells were harvested for analysis or differentiated into osteoblasts for 21 days as above prior to analysis.

### SOX11 knockdown

Lentiviral vectors TRC1-pLK0.1 containing a non-target, scrambled (NT) shRNA (5’-GCGATAGCGCTAATAATTT-3’ SHC202; Sigma-Aldrich) or a shRNA specific for human SOX11 (100% identity to the equine SOX11 sequence) (5’-CTGGTGGATAAGGATTTGGAT-3’ clone NM_003108.3-1235s1c1; Sigma-Aldrich) were created by transfection into HEK293T packaging cells. As a positive control to visualise successful transfection and transduction, TRC2-pLKO.5-puro-CMV-TurboGFP plasmid was used (SHC 203; Sigma-Aldrich). HEK293T cells were plated at 1 x10^5^ cells per well of a 6 well plate and transfected with 1 µg of TRC2-pLK0.1 plasmid, 750 ng psPAX2 and 250 ng pMD2.G. pMD2.G and psPAX2 were a gift from Didier Trono (Addgene plasmids #12259 and 12260; http://n2t.net/addgene:12259 http://n2t.net/addgene:12260; RRID:Addgene_12259 and 12260). Packaging cell supernatant was collected after 48 h, filtered through a 0.45 µm filter (Millipore, Billerica, US) and used immediately. Skin fibroblasts (from three Thoroughbreds and one Welsh Mountain pony) were seeded at 2×10^5^ cells per well of a 6 well plate 24 h before infection. To achieve 80% of cells transduced (determined via GFP expression), up to three rounds of infection were performed in the presence of 10 µg/ml polybrene (Sigma-Aldrich). 48 h after the last round of infection, puromycin (Sigma-Aldrich) selection was carried out as described above and then cells were used for RNA extraction.

### ELISA

Whole cell protein extract was prepared from undifferentiated skin cells (overexpressing SOX11 or control) using three rounds of freeze-thawing in extraction buffer containing 20 mM Hepes pH7.9, 450 mM NaCl, 0.4 mM EDTA, 25% glycerol, 1 mM PMSF and the supernatant collected by centrifugation. An equine specific COL3A1 ELISA (EKU11662, Biomatik, Delaware, USA) was subsequently used to measure the amount of COL3A1 in 300 ng of total whole cell extract. Measurements were made on a microplate absorbance reader (Mplex (Infinite M200); Tecan, Switzerland).

### Electrophoretic mobility shift assay (EMSA)

Nuclear protein extract (NPE) was extracted from skin fibroblasts derived from two different Welsh Mountain ponies that were overexpressing human SOX11 (as described above) using NE-PER Nuclear and Cytoplasmic Extraction Reagents (Thermofisher) according to the manufacturer’s instructions. EMSAs were carried out in accordance with the protocol of LightShift chemiluminescent EMSA kit (Thermofisher). For this purpose, 20 fmol of biotinylated double-stranded oligonucleotides: 5’-biotin-GAATCACTCAAATATT<u>T</u>GCCTGTCAGGTCC −3’ or 5’-biotin-GAATCACTCAAATATT<u>G</u>GCCTGTCAGGTCC −3’, were incubated in 20 µl of reaction mixture (10x binding buffer, 2.5% (v/v) glycerol, 5 mM MgCl_2_, 50 ng/µl poly (dI·dC), 0.05% NP-40, 50mM KCl, 10mM EDTA) at room temperature for 20 min. For the shift assay, labelled probes were incubated with a total of 30 µg of NPE. For the competitor assay the nuclear extract was pre-incubated with 200-fold molar excess of unlabelled probes before adding the labelled probe. For the supershift assay, the nuclear extract was incubated, for 1 h at room temperature, with 5.8 µg of rabbit anti-SOX11 (sab4200450) (Sigma) after adding labelled probe. 5 µl of loading buffer was then added to stop the reaction. The samples were loaded onto Invitrogen™ DNA Retardation Gels (6%) (ThermoFisher) and run in 0.5X TBE for 1 h at 100 V. The gel was transferred to Invitrogen™ BrightStar™-Plus Positively Charged Nylon Membrane (Thermofisher) for 30 min at 380 mA. DNA membrane was cross-linked for 10-15 min on a transilluminator equipped with 312nm UV-bulbs. The detection was performed using LightShift chemiluminescent EMSA kit (Thermofisher) according to manufacturer’s instructions. Technical controls for the assay were carried out using Epstein-Barr Nuclear Antigen probes and protein extracts as provided in the kit.

### Construction of the COL3A1-5’UTR luciferase reporter plasmids

The reference allele (TT) COL3A1-5’ UTR fragment (ECA18: 65598913-65602132, EquCab 3.0) was PCR-amplified from genomic homozygous TT allele Thoroughbred DNA using the primers: Froward 5’-ctattggtaccGAGGAATTGGTCACAGGAATCAC-3’ and Reverse 5’-ctggaagcttGCAGTTCAAAGTAGCACCATC-3’, showing restriction sites for KpnI and HindIII, respectively. The alternative allele (GG) COL3A1-5’ UTR fragment was created from the reference allele (TT) PCR product using the Q5® Site-Directed Mutagenesis Kit (New England BioLabs, Massachusetts, USA). The sequence of primers carrying out the alternative allele: Forward 5’-CTCAAATATTGGCCTGTCAGG-3’ and Reverse 5’-TGATTCCTGTGACCAATTC-3’, were designed using online NEB primer design software, NEBaseChanger™. pGL4.10[Luc2] (#E6651 Promega, Southampton, UK) and PCR products were digested with KpnI and HindIII, and column-purified. The KpnI-HindIII fragments with the reference or alternative allele were then cloned into the previously digested pGL4.10[Luc2] vector. In each step, PCRs were performed by Q5 High-fidelity DNA Polymerase (New England Biolabs) and PCR products were verified by gel electrophoresis and sequencing (performed by DNA Sequencing and Service, University of Dundee, UK). In each case, ligation products were transformed in DH5α competent cells (ThermoFisher), which were then plated onto ampicillin-containing (100 µg/ml) LB agar (Sigma-Aldrich), and individual colonies were analyzed by restriction enzyme digest and sequencing after 24 h of growth at 37 °C.

### Luciferase assays

Skin fibroblasts from six Thoroughbreds (0.3 x 10^5^) with a range of overall risk scores were transfected in 24-well plates using Lipofectamine3000 DNA transfection reagent (ThermoFisher). The cells in each well were co-transfected with 1.4 µg of reference allele (TT) or alternative allele (GG) COL3A1-5’UTR firefly luciferase reporter vector and 0.14 µg of internal control Renilla luciferase vector, pGL4.74[hRluc/TK] (Promega), as the control for transfection efficiency. pGL4.10[Luc2] vector with no insertion was used as a promoter-less negative control. 48 h after transfection, cells were harvested and assayed using Dual-Glo® luciferase assay system (Promega). The luciferase activities were measured using luminescence reader (Mplex (Infinite M200); Tecan). Promoter activity was expressed as relative firefly luciferase activity after normalization to Renilla luciferase activity.

### Statistical analysis

Statistical analysis of qPCR and luciferase data was performed using SPSS. Shapiro Wilks normality testing was performed to confirm a normal distribution of the data (raw or log transformed data) and Levene’s test used to confirm equal variance between the groups. For comparisons of two groups the Student’s t-test (unpaired, two tailed) was used, for more than two groups ANOVA was used with a Tukey’s post hoc test. A Chi-square test was used to measure the association of the SNP with fracture. In all statistical tests a significance threshold of p<0.05 was set.

## Results

### Skin fibroblasts can differentiate into osteoblasts in vitro

Skin fibroblasts were isolated from 44 Thoroughbred horses, DNA extracted and cells banked. Genotyping of the samples was performed to calculate their relative polygenic risk score for fracture. We then selected the three male samples with the lowest genetic risk and the three male samples with the highest genetic risk for use in osteoblast differentiation. These samples were at the ends of the risk spectrum when compared to the scores from 382 Thoroughbreds representing the general population (Figure 2).

**Figure 2.**
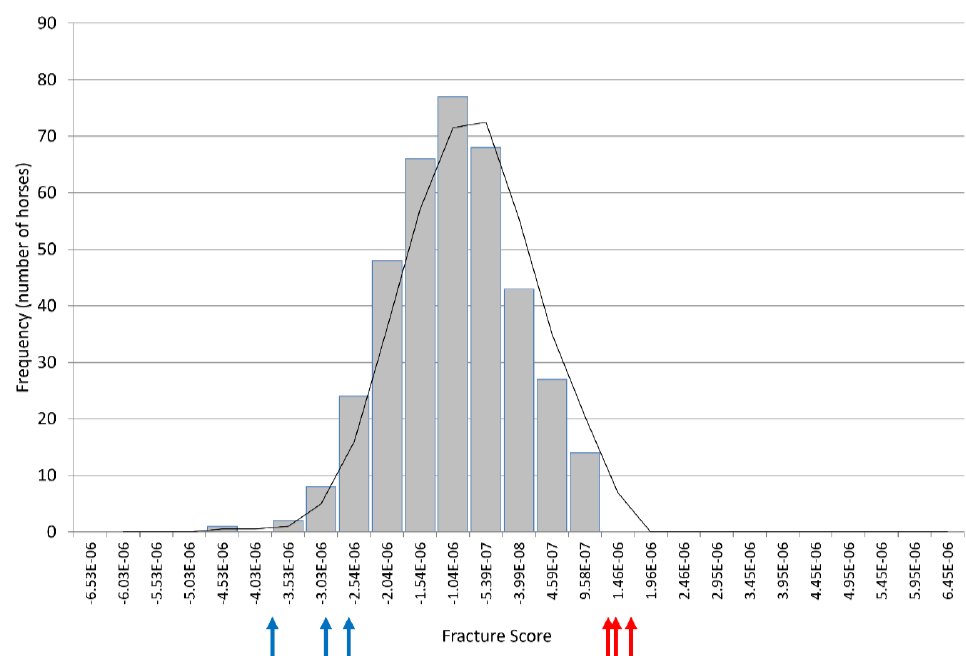
The polygenic risk scores of 382 Thoroughbreds of unknown fracture status (representing the general population) with arrows indicating the scores for the skin cells selected as the three highest (red) and three lowest (blue) risk.

Osteoblast differentiation was carried out at using two concentrations of β-glycerophosphate, 2 mM and 10 mM. We demonstrated that after 21 days of culture the skin fibroblasts expressed a range of different osteoblast associated genes and this was not significantly affected by the level of β-glycerophosphate (Figure 3A). Furthermore, both 2 mM and 10 mM β-glycerophosphate result in the production of a mineralised matrix (Figure 3B).

**Figure 3.**
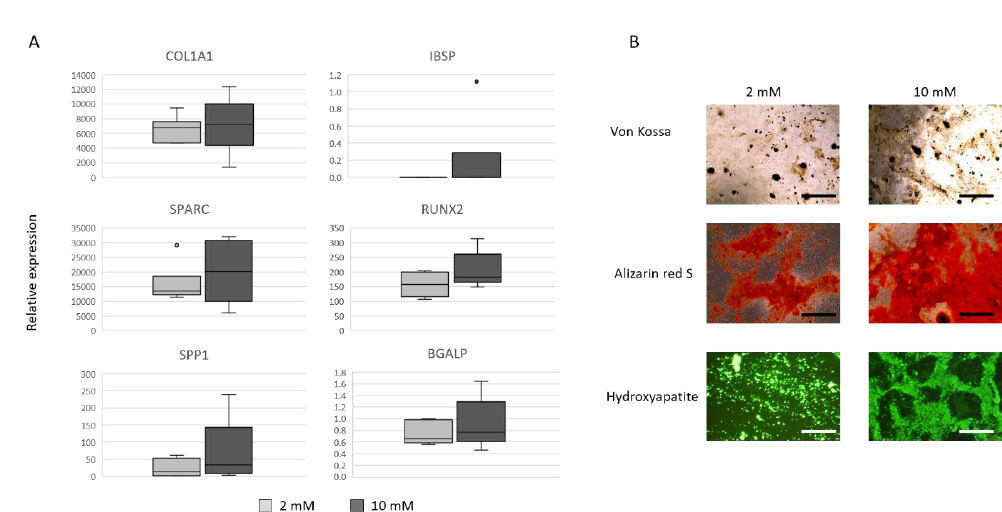
Skin fibroblasts can differentiate into mineralising osteoblasts. A) Box and whisker plots showing osteoblast associated gene expression in skin fibroblasts from six individual horses. There are no significant differences in the expression of osteoblast associated genes following 21 days of differentiation in media with either 2 mM (light grey bars) or 10 mM β-glycerophosphate (dark grey bars). Expression is shown relative to the housekeeping gene. B) A mineralised matrix is deposited following osteoblast differentiation in either 2 mM or 10 mM β-glycerophosphate. Positive staining for calcium deposition is shown in black for von Kossa and red for Alizarin Red S. Positive hydroxyapatite staining detected under a fluorescent light is shown in green. Scale bars = 100 µm. Images are representative of six biological replicates using cells at passage 5-6.

### Candidate gene expression analysis reveals a significant difference in STAT1 and COL3A1 gene expression in skin-derived osteoblasts from high and low risk horses

Previous work had demonstrated that a region on ECA18 was significantly associated with fracture risk (Blott *et al*. 2014). This region contains 27 protein coding genes with a known function. Of these, 11 have previously published associations with bone formation or fracture, as described in the introduction. We therefore measured the expression of these 11 candidate genes in skin-derived osteoblasts from the three horses at highest and three horses at lowest genetic risk of fracture (as shown in Figure 2). *STAT1* was expressed at significantly higher levels in cells derived from horses at high risk of fracture than in horses at low risk of fracture, and *COL3A1* was expressed at significantly higher levels in cells derived from horses at low risk of fracture (Figure 4). No expression of *ZNF804A* was detected, and no significant differences in the expression of the other genes between high and low risk horses were found.

**Figure 4.**
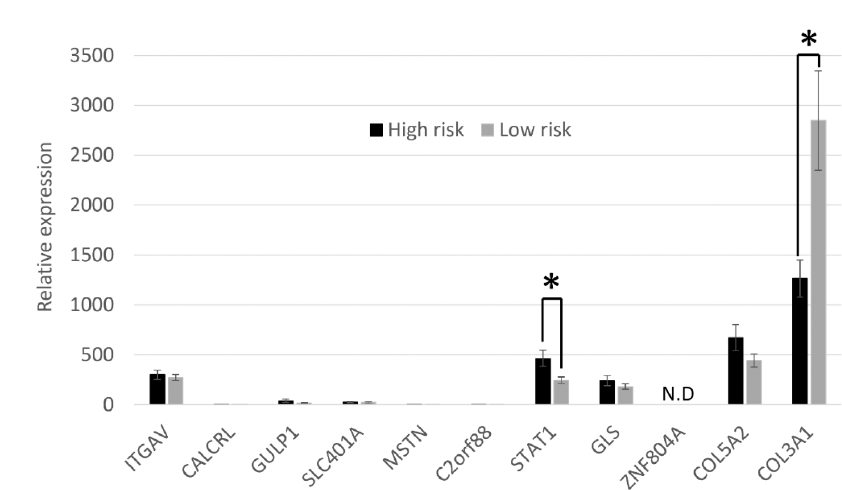
Expression of candidate genes from the fracture associated region on ECA18 in skin cell derived osteoblasts. There is a significant difference in the expression of *STAT1* and *COL3A1* between cells derived from high and low risk Thoroughbred horses. Expression is shown relative to the housekeeping gene. Error bars represent the s.e.m of cells derived from three individual horses, each differentiated into osteoblasts duplicate (once in 2 mM β-glycerophosphate and once in 10 mM β-glycerophosphate). *p<0.05. N.D = expression not detected. Cells were at passage 5-6. STAT1 can sequester the essential regulator of osteoblast differentiation RUNX2 in the cytoplasm of cells. We therefore examined the cellular localisation of STAT1 and RUNX2 in either the cells derived from high and low risk horses but no significant differences were observed (Figure 5).

**Figure 5.**
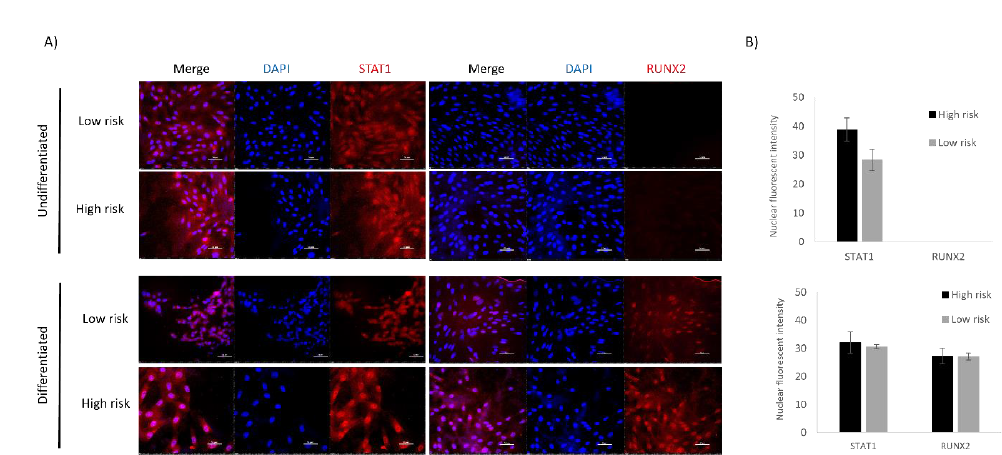
STAT1 and RUNX2 have similar cellular localisations in cells derived from high risk and low risk horses. A) Immunocytochemistry demonstrates STAT1 (red) expression in undifferentiated skin fibroblasts and following osteoblast differentiation, but RUNX2 (red) is only present following differentiation. DAPI staining of the nucleus is shown in blue. Scale bar = 50 µm. Images are representative of replicates using cells derived from three different donors. B) Quantification of the intenisity of the staining in the nuclei shows no significant differences between cells derived from high and low risk horses. Error bars represent the s.e.m from cells derived from three different donors.

### WGS revealed a SNP in the region upstream of COL3A1 that is predicted to result in the loss of a Sox11 binding site and is significantly associated with fracture risk

Preliminary whole genome sequencing on two cases and two control did not reveal any exonic SNPs in the associated region on chromosome 18 that were predicted to affect protein structure or function. However, there were a large number of intronic/intergenic variants. One variant was located 3154 bp upstream of the transcriptional start site (TSS) of *COL3A1* in (position ECA18:65,598,945 T>G EquCab 3.0). This region is highly conserved between human and horse and ENCODE data demonstrates that the human region has markers of an enhancer/promoter (e.g. multiple transcription factor binding sites and histone H3 lysine 4 methylation, H3K4me). The Functional Annotation of Animal Genomes (FAANG) database also demonstrates the presence of H3K4me and H3K27ac (histone H3 lysine 27 acetylation) in the region covering the SNP in adipose tissue, ovary and skin. The Genomic Evolutionary Rate Profiling (GERP) conservation score (Cooper *et al*. 2005) of the SNP is −0.28. The minor allele frequency (MAF) was 0.32 in our 91 case horses and 0.22 in our 86 controls (i.e. 0.27 on average). Across 25 different breeds of horse (Jagannathan *et al*. 2019), the MAF was 0.13. In many breeds, the MAF was very low, whereas in US/European Thoroughbreds it was 0.53, similar to Japanese Thoroughbreds (0.5). It was also very high in Quarter horses (0.66) (Supplementary table 3). The genotype of ten Thoroughbreds with heterozygous or homozygous genotypes was confirmed to be accurate using the Integrative Genome Viewer.

The SNP is predicted to change a transcription factor binding motif from a SOX9/10/11 binding site to a KLF13 binding site (Figure 6A). Allelic discrimination assays were performed (Figure 6B and C) and demonstrated that the G variant is significantly associated with fracture (p<0.01).

**Figure 6.**
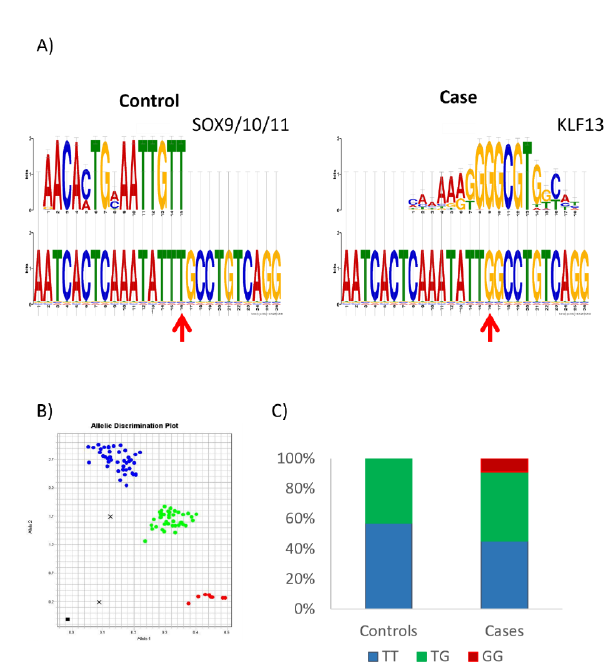
An associated SNP of interest lies upstream of COL3A1. A) A SNP identified through preliminary whole genome sequencing is predicted to cause the loss of a SOX9/10/11 binding site and the gain of a KLF13 binding site. Red arrow indicates the position of the SNP. Analysis output from TomTom. B) Representative allelic discrimination plot of the *COL3A1* upstream SNP. Blue indicates TT homozygotes, green indicates TG heterozygotes, red indicates GG homozygotes.C) The percentage distribution of the genotypes in 86 horses without a fracture (controls) and 91 horses with catastrophic fractures (cases).

### SOX11 is expressed in undifferentiated and osteoblast-differentiated skin fibroblasts, but is not differentially expressed between high and low risk horses

SOX9 and 10 are both members of the SOXE group and have overlapping functions (Haseeb & Lefebvre 2019). SOX9 is a master regulator of cartilage differentiation and is usually decreased in osteoblast differentiation (Haseeb *et al*. 2021). In contrast SOX11, member of the SOXC group (Dy *et al*. 2008), is required for osteoblast precursor survival, proliferation and differentiation (Gadi *et al*. 2013). We demonstrated that *SOX11* is expressed in skin fibroblasts before and after differentiation into osteoblasts, but there is no significant difference in *SOX11* expression levels between high and low risk horses (Figure 7).

**Figure 7.**
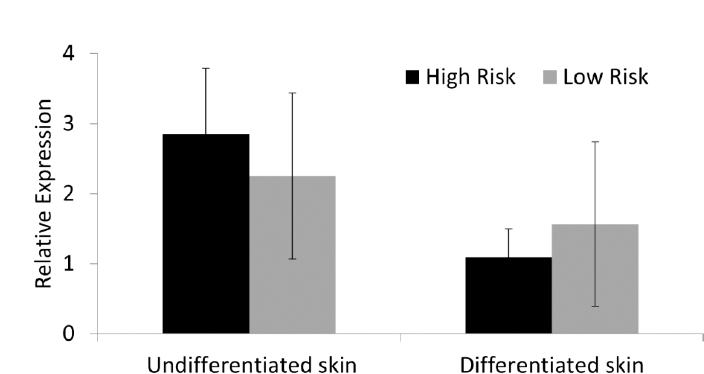
*SOX11* expression. *SOX11* is expressed in Thoroughbred undifferentiated skin fibroblasts and osteoblasts derived from skin fibroblasts. There are no significant differences in expression between cells derived from high and low risk horses. Expression is shown relative to the housekeeping gene. Error bars represent the s.e.m of cells derived from five individual horses. Cells were at passage 2-6.

### SOX11 modulation in skin fibroblasts results in significant changes in the expression of COL3A1

Skin fibroblasts were stably transfected to overexpress the human *SOX11* gene. In the undifferentiated skin fibroblasts, this resulted in a large, significant increase in *SOX11* expression (Figure 8Ai) and a significant increase in *COL3A1* expression (Figure 8Aii). This suggests that SOX11 regulates *COL3A1*. Following osteoblast differentiation of the skin fibroblasts, there was no significant difference in the expression of *COL3A1* in cells overexpressing *SOX11* compared to the control cells. This is because the relative level of *SOX11* overexpression was reduced after differentiation which may reflect a lower activity of the CMV promoter (used to drive overexpression) in osteoblasts (Figure 8A). No differences were observed in the total protein level or cellular localisation of COL3A1 in undifferentiated skin fibroblasts following SOX11 overexpression (Supplementary Figure 2). However, lentiviral transfection of undifferentiated skin fibroblasts to express an shRNA to *SOX11* produced a significant decrease in *COL3A1* (Figure 8B). Taken together this data demonstrates that SOX11 can regulate *COL3A1* expression.

**Figure 8.**
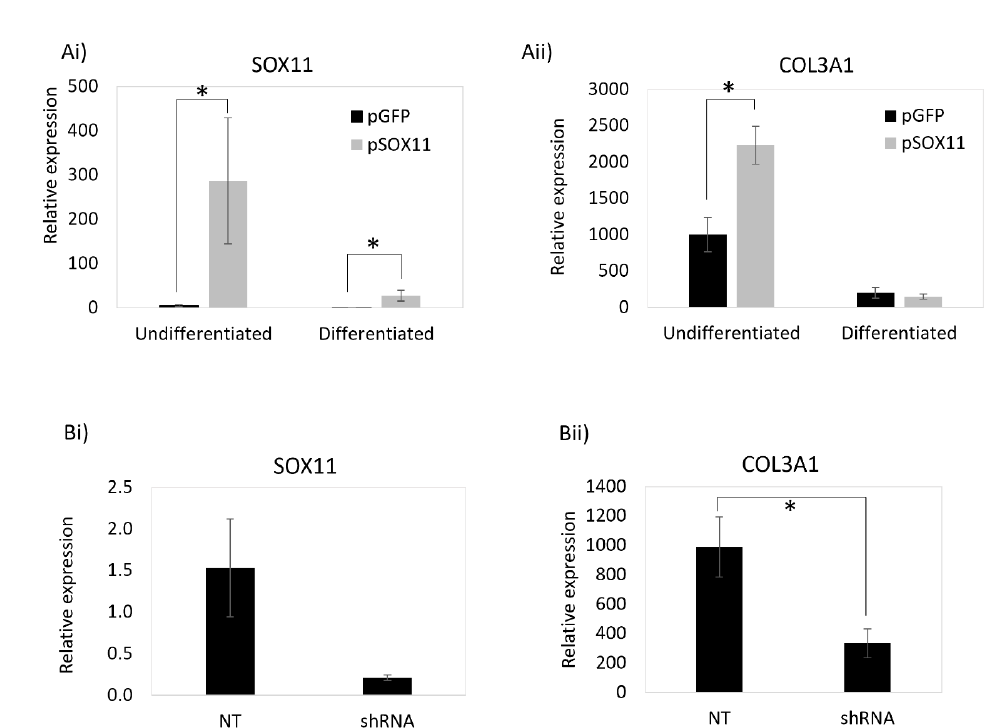
SOX11 stable modulation results in significant changes in *COL3A1* expression. Ai) Stable transfection of a *SOX11* overexpression plasmid (pSOX11) results in a significant increase in *SOX11* expression in skin fibroblasts compared to control cells (pGFP). Aii) This results in a significant increase in *COL3A1* expression in the undifferentiated cells. B) Lentiviral expression of a shRNA to *SOX11* reduces *SOX11* (Bi) and *COL3A1* (Bii) expression in undifferentiated skin cells. Expression is shown relative to the housekeeping gene. Error bars represent the s.e.m from cells derived from three or four different donors. *p<0.05. Cells between passages 5 and 10 were used in these experiments.

### The region containing the SNP binds to nuclear protein from equine cells

EMSA was performed using nuclear extracts from undifferentiated skin fibroblasts overexpressing SOX11. We observed a gel shift indicating protein binding to the probe whether the probe was containing the TT control allele or the fracture associated GG allele. However, the addition of a SOX11 antibody did not result in a supershift (Figure 9).

**Figure 9.**
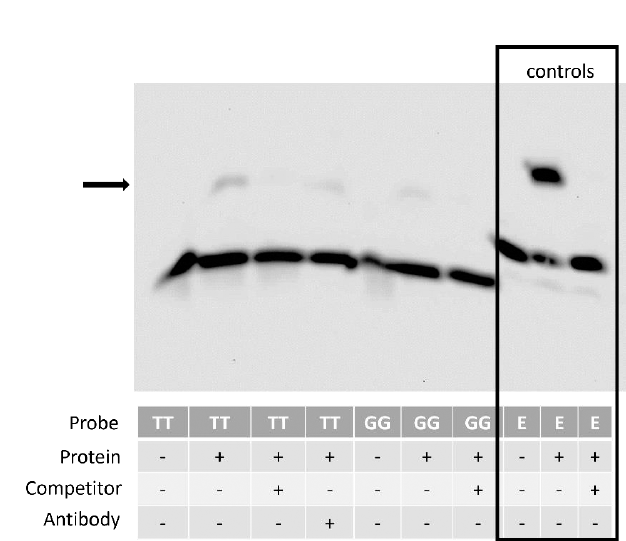
The region upstream of COL3A1 containing the SNP can bind to a nuclear protein. EMSA revealed that a probe containing either the wild-type (TT) or variant (GG) allele was capable of binding to a nuclear protein as seen by the shift (arrow). The use of an unlabelled competitor probe prevented this shift from occurring. The addition of an antibody to SOX11 did not result in a detectable supershift. Image representative of results observed using nuclear extracts from cells isolated from two different horses. The boxed area represents a technical control for the assay, carried out using a biotin labelled probe for Epstein-Barr Nuclear Antigen (EBNA, labelled E in the figure), an unlabelled probe and EBNA protein extract. The image is of the entire full size blot and has not been cropped.

### The region containing the SNP has promoter activity and the SNP affects reporter gene expression in a genetic background dependent manner

We carried out luciferase assays in undifferentiated skin fibroblasts and demonstrated that the region upstream of the equine *COL3A1* gene including the SNP does have promoter activity (Figure 10). However, the effect of the SNP is variable and depends on the genetic background of the individual. In skin fibroblasts derived from some horses, the presence of the SNP caused a reduction of the promoter activity (Figure 10 A and B). In contrast in other horses, the presence of the SNP caused an increase in promoter activity (Figure 10 C and D) and, in some horses, the presence of the SNP had no significant effect on promoter activity (Figure 10 E and F). There was no correlation between the result and the overall risk score of the horse from which the cells were derived. Each assay was carried out on three to six technical replicates performed simultaneously. However, additional replicates were also performed using the same donor horse but on replicates set up at three different times. For the same donor cell line, the results were consistent between these independent experiments (supplementary Figure 3).

**Figure 10.**
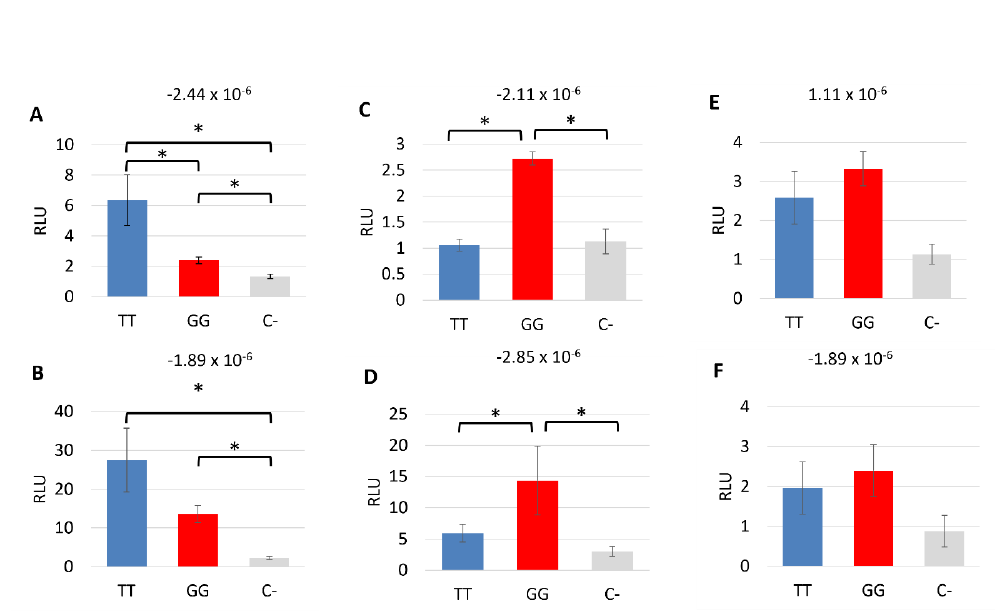
The region upstream of COL3A1 containing the SNP has promoter activity and the effect of the SNP varies with the donor cells used. Luciferase assays showing the relative light unit (RLU) for the 3220 bp region upstream of COL3A1 containing either the reference allele (TT) or the alternative allele (GG) compared to a promoterless control (C-). Error bars represent the s.e.m of three to six technical replicates. Graphs A-F represent the assay being performed in skin fibroblasts from six different donor horses, the risk score for each horse is displayed above each graph. *p<0.05.

We were not able to perform the luciferase assays in the skin cells following osteogenic differentiation as the cells cannot be transfected (data not shown).

## Discussion

In this study we have employed polygenic risk scores using genome wide information to facilitate the establishment of an *in vitro* system to study bone gene regulation in horses at high and low genetic risk of fracture. These samples were not part of the original cohort of cases and controls used to derive the risk score. Validating the polygenic risk score in a new cohort of cases and controls is currently ongoing.

Skin fibroblasts were selected as the starting cell type due to their accessibility, and ease of culture, which allowed us to create a bank of cells from a large number of horses representing the general Thoroughbred population and select cells which were at very high or very low genetic risk. Primary osteoblasts could not be used as they have very limited proliferative capacity *in vitro* (Kassem *et al*. 1997). Skin fibroblasts have previously been shown to share properties with MSCs *in vitro* and can be driven to differentiate into mineralising osteoblasts (Lorenz *et al*. 2008; Halcsik *et al*. 2013). We confirm this finding using horse skin fibroblasts, although in future studies it may be possible to use other cell types such as primary MSCs (Glynn *et al*. 2013) or induced pluripotent stem cells (iPSCs) (Nagy *et al*. 2011; Sharma *et al*. 2014; Bavin *et al*. 2015) as a starting cell type. We selected skin fibroblasts only from male Thoroughbreds as some previous studies have shown an association of catastrophic fracture with sex (Anthenill *et al*. 2007; Georgopoulos & Parkin 2017; Sun *et al*. 2019), but this is future studies using larger numbers of female and male cells would be warranted. Our cell-based system was established to allow us to control for environmental variability, but it is well known that other factors can influence cell behaviour *in vitro* e.g. donor age. This study only used cells from young Thoroughbred horses between the ages of two and four years old, but we cannot rule out that other biases may have existed in the cells which affected the results.

To differentiate the skin fibroblasts into osteoblasts we tested two concentrations of β-glycerophosphate. 10 mM β-glycerophosphate has been commonly applied to equine induced pluripotent stem cells (iPSCs) (Baird *et al*. 2018), equine MSCs (Guest *et al*. 2008; Radcliffe *et al*. 2010; Burk *et al*. 2013) and human and mouse skin fibroblasts (Lorenz *et al*. 2008; Halcsik *et al*. 2013). However, there have been reports that levels of β-glycerophosphate over 2 mM can lead to non-physiological mineral deposition (Chung *et al*. 1992). In this study we saw no significant effect of β-glycerophosphate concentration on gene expression or matrix mineralisation, but additional biological replicates may enable the detection of smaller differences in differentiation.

We used the skin-derived osteoblasts to carry out a candidate gene expression analysis of genes that lie within a region on ECA18 that has previously been shown to be associated with catastrophic fracture (Blott *et al*. 2014). All of the candidate genes were expressed in the skin-derived osteoblasts with the exception of *ZNF804A*, which was previously suggested to be a candidate gene for fracture risk (Blott *et al*. 2014) but was not expressed in our experimental system. Of the candidate genes examined, *COL3A1* and *STAT1* were differentially expressed in the high and low risk cells. In the future, analyses using RNA sequencing to examine global gene expression profiles would be beneficial. As we had a limited number of samples (three high risk and three low risk) future work using larger cohorts may detect other differentially expressed genes. However, we were limited in the samples we had available as we wanted to ensure that we selected cells that were at the ends of the risk spectrum (Dobrindt *et al*. 2021).

STAT1^-/-^ mice have a significant increase in bone mineral density, mineral content and bone growth (Xiao *et al*. 2004). This is due to increased osteoblast differentiation (Kim *et al*. 2003). STAT1 acts to sequester the essential regulator of osteoblast differentiation RUNX2 in the cytoplasm of cells. Therefore, in the absence of STAT1, more RUNX2 enters the nucleus to activate osteoblast-associated gene transcription (Kim *et al*. 2003). Furthermore, it has been shown in mice that STAT1 inhibition accelerates fracture healing (Tajima *et al*. 2010). The control of RUNX2 expression and activity needs to be tightly controlled, as over-expression of *Runx2* leads to mice that have osteopenia and suffer from multiple fractures (Liu *et al*. 2001). Therefore in our study, the higher level of *STAT1* observed in the high risk cells may contribute to mis-regulation of other bone specific genes and predisposition to fracture. However, we were unable to demonstrate any differences in the cellular localisation of STAT1 or RUNX2 in the high and low risk horses. Furthermore, our WGS was not able to detect any putative, functional SNPs in the vicinity of the *STAT1* gene. This may reflect the small number of WGS samples used. In the future it would be beneficial to perform WGS on more samples to increase our power to detect SNPs and/or other variants (e.g. structural variants). However, our results may also suggest that other factors are regulating expression of *STAT1*, such as long-range enhancers. Further work to determine the mechanisms behind the differential expression of *STAT1* and identify any downstream effects are therefore required.

COL3A1^-/-^ mice are not viable. However, COL3A1^-/+^ mice have reduced bone volume, bone density and trabecular thickness and COL3A1^-/-^ MSCs have reduced osteoblast differentiation compared to wild type MSCs (Volk *et al*. 2014). In humans loss of function mutations of *COL3A1* result in Ehlers Danlos Syndrome and these patients have a tendency to fracture, low bone mass and abnormal bone structures (Yen *et al*. 2006). Therefore, in our study, the lower levels of *COL3A1* in the cells derived from horses at high risk of fracture, may contribute to their risk. *COL3A1* regulation is not well defined, but our WGS data revealed a SNP 3154 bp upstream of the *COL3A1* transcriptional start site. As our WGS was performed on only two cases and two controls we carried out allelic discrimination genotyping in a further 86 controls and 91 cases and found that the SNP was significantly associated with fracture.

The SNP exists in a region which contains histone modifications in multiple equine tissues suggesting promoter/enhancer function. No histone modifications were found in the FAANG database for bone tissue but these tissues had lower quality metrics and the data is more limited than for other tissues (Kingsley *et al*. 2021). Although its GERP score does not indicate high levels of evolutionary conservation, there are limitations to using this approach to identify deleterious mutations, particularly in noncoding sites (Huber *et al*. 2020). The MAF of the SNP varied between breeds, with some breeds showing very low frequencies and Thoroughbreds and Quarter horses showing much higher frequencies. Interestingly, the MAF appeared to be lower in our UK Thoroughbreds compared to variant catalogues of international Thoroughbreds which may reflect some population differences. The higher allele frequencies in some Thoroughbred populations and the small difference in the cases versus the controls in the UK population (0.32 versus 0.22) suggest it is unlikely to be a variant of high effect. However, it may still contribute to fracture risk, which is a complex disease and therefore most likely explained by multiple variants of small to moderate effect size.

Bioinformatics tools suggested that the SNP may decrease SOX11 binding and promote KLF13 binding. KLF13 is a transcription factor involved in erythropoiesis (Gordon *et al*. 2008). It has also been shown to act as a tumour suppressor in colorectal cancer cells (Yao *et al*. 2020) and to be a novel regulator of heart development (Lavallée *et al*. 2006). It is not clear if KLF13 has any role in bone formation or function, but it has been shown to be upregulated by dexamethasone during osteoblast differentiation (Leclerc *et al*. 2004) and is expressed in the developing mouse skeleton (Martin *et al*. 2001). The potential effect of the SNP on KLF13 binding and KLF13 regulation of *COL3A1* therefore warrants future investigation.

Although SOX11 is involved in bone differentiation (Gadi *et al*. 2013; Xu *et al*. 2015), it has not previously been shown to regulate *COL3A1*. In skin fibroblasts the stable knockdown of *SOX11* through an integrated shRNA resulted in a significant decrease of *COL3A1* expression. Likewise, overexpression of SOX11 produced a significant increase in *COL3A1* gene expression. This was not reflected at the protein level, however protein and RNA levels do not always correlate, particularly in response to perturbation (Vogel & Marcotte 2012). This may suggest that COL3A1 is also regulated at the level of translation. Following differentiation, no significant effect of SOX11 over-expression was observed but this likely reflects the lower levels of *SOX11* over-expression following differentiation. Promoter activity has been correlated with cell type, and CMV in particular shows variable expression in different cells (Qin *et al*. 2010). Future work to test the activity of different promoters in equine osteoblast cells is required to overcome this limitation.

The promoter region of the equine *COL3A1* gene has not previously been defined. EMSA using equine nuclear extracts from SOX11 overexpressing cells demonstrated protein binding to the region of DNA containing the SNP. Protein binding was observed using DNA containing both the control (TT) and alternative (GG) allele. Fainter binding was observed in the presence of the GG allele, but we cannot rule out the possibility that this reflects technical variation in the assay, rather than a true reduction in binding. We were unable to detect a supershift using an antibody to SOX11 and are therefore unable to confirm the identity of the binding proteins. This may reflect a genuine lack of SOX11 binding to our region of interest or it may reflect a lack of detection. SOX11 has previously been reported to contain a conserved domain which inhibits DNA binding *in vitro* and prevent detectable binding using EMSAs *(Wiebe et al. 2003; Dy et al. 2008)*, although other studies have detected SOX11 binding using *in vitro* translated protein (Kan *et al*. 2013). We used nuclear extracts from SOX11 overexpressing cells to ensure that robust levels of SOX11 were present. However, the SOX11 protein is tagged with GFP. This may interfere with SOX11 antibody binding and may account for the lack of supershift. Alternatively, it may reflect the weak binding of SOX11 under the conditions used in this *in vitro* assay. The GG allele is predicted to result in both the loss of the SOX11 binding site and gain of a KLF13 binding site. As both proteins are of a similar size, it is not possible to determine if the binding proteins to the TT probe are the same as those binding to the GG probe without further analysis. It is also possible that different results may be obtained if we were to carry out the EMSA using nuclear extracts from cells differentiated into osteoblasts.

The SOX11 knockdown and overexpression assays were performed using cells from both Thoroughbreds and Welsh Mountain ponies and produced similar results. The EMSA was performed using cells from Welsh Mountain ponies. This was done to protect our stocks of Thoroughbred skin cells, which do not proliferate indefinitely in culture. However, these experiments should be independent of the breed of horse used as they are not looking at the effect of genetic variants, but demonstrate only that SOX11 can regulate COL3A1 expression and that the region of interest can bind to nuclear proteins. Such experiments that are usually performed in laboratory animal cells to understand human biology (Gadi *et al*. 2013; Xu *et al*. 2015) given that gene functions are well conserved between species.

Luciferase assays in undifferentiated skin cells demonstrated that the proximal region (from −3186 to +34) of *COL3A1* containing the SNP of interest does have promoter activity. We attempted to perform the luciferase assays in an osteoblast cell type, but it was not possible to transfect the skin cells following osteoblast differentiation, possibly due to the presence of the mineralised matrix they produce (data not shown). To our knowledge, there are no reports on successful transfection of cells following osteogenic culture and mineralisation. There are also very few reports on the successful culture of primary equine osteoblasts and we did not have access to suitable tissue samples to derive these cells. Attempts to use primary equine osteoblasts for these experiments were therefore outside the scope of this paper.

Furthermore, the luciferase assays demonstrated that the effect of the SNP was dependent on the genetic background of the horse, with different effects observed using cells from different horses. This reflects the complexity of studying complex genetic conditions even in an *in vitro* system where environmental effects are minimised. The results suggest that multiple genetic modifications may interplay to regulate *COL3A1* expression and further work to identify other pathways that may converge on COL3A1 is required.

The major limitation of this study was the small sample sizes as described. In addition, the luciferase assays, EMSAs and SOX11 modulation experiments were only successfully performed in the undifferentiated fibroblasts due to the technical limitations described. However, this work is the first demonstration that SOX11 is capable of regulating *COL3A1* and demonstrates that a specific region upstream of *COL3A1* has promoter activity. Furthermore, the fibroblasts used have osteogenic differentiation potential. Bone remodelling and fracture repair involves the activities of both mature osteoblasts and progenitor cells (Bragdon & Bahney 2018) and future work to understand the regulation and role of COL3A1 in both cell types during bone remodelling is warranted.

In conclusion, we have established a cell-based system to allow us to study the genetic basis of fracture in Thoroughbred horses. In support of previous work identifying a fracture associated region on ECA18 (Blott *et al*. 2014; Tozaki *et al*. 2020), we demonstrated that one of the genes within this region, *COL3A1,* is differentially expressed in cells from horses at high and low risk of fracture and that this may in part be due to the presence of an associated SNP in the *COL3A1* promoter. The SNP may affect SOX11 binding, and we demonstrate that SOX11 modulates *COL3A1* expression. These findings support future work using this system to identify other genes that are differently expressed across the genome and other SNPs that are involved in fracture risk. This approach has the potential to be very valuable in teasing apart the complexity of the genetic basis for fracture risk.

## Statements and Declarations

### Compliance with Ethical Standards

Equine skin fibroblasts were taken from horses that had been euthanised for reasons unrelated to this study and with the consent of the Animal Health Trust Ethical Review Committee (AHT_02_2012) and Royal Veterinary College Clinical Research Ethical Review Board (URN 2021 2035-2).

### Availability of data and materials

All data generated and analysed during this study are included in this published article and its supplementary information files. The WGS data have been deposited in the European Nucleotide Archive (ENA) at EMBL-EBI under accession number PRJEB60529. (https://www.ebi.ac.uk/ena/browser/view/PRJEB60529).

### Author contributions

D.J.G. conceived and designed the study. E.P.L. performed the majority of the experimental work, A.B. carried out the WGS analysis, skin fibroblast banking and polygenic risk scoring, R.M. performed the overexpression work and A.C.R performed the immunocytochemistry and analysis for STAT1 and RUNX2. S.B. designed the polygenic risk scoring protocol. S.A.D-A performed the minor allele frequency analysis. D.J.G. supervised the project, secured the funding and drafted the manuscript. All authors approved the manuscript.

## Acknowledgements

This study was kindly funded by the Horserace Betting Levy Board (vet/prj/792). A.B. was funded by the Anne Duchess of Westminster Charitable Trust who also funded the whole genome sequencing. The Alborada Trust fund A.C.R.

