## Supplementary data for "A functional single nucleotide polymorphism upstream of the collagen type III gene may contribute to catastrophic fracture risk in Thoroughbred horses"

**Supplemental Material**

| **Gene** | **Forward primer** | **Reverse primer** |
| --- | --- | --- |
| 18S RNA | CCCAGTGAGAATGCCCTCTA | TGGCTGAGCAAGGTGTTATG |
| GAPDH | CGACCACTTTGTCAAGCTCA | GTCCACCACCCTATTGCTGT |
| ACTB | CCAGCACGATGAAGATCAAG | GTGGACAATGAGGCCAGAAT |
| COL1A1 | TGCGAAGACACCAAGAACTG | GACTCCTGTGGTTTGGTCGT |
| SPARC | TGGCGAGTTTGAGAAGGTGT | TTTGCAAGGCCCGATGTAGT |
| SPP1 | AGCCCCAGGAAAAATCGCTG | GGCATAAGCAAATCACGGCA |
| IBSP | GGACTGCACACGGAAACAATC | ACAGGCCATTCCCAAAATGC |
| RUNX2 | CCAAGTGGCAAGGTTCAACG | AACTCTTGCCTCGTCCACTC |
| BGALP | GTCTCGGGGTTCCAAGGTTA | AATCTCTGGTAGCTGTGTTGGT |
| ITGAV | GATGCTGATGGACAGGGATT | AAACTACCAGGACCACCAAGAA |
| CALCRL | TTTGTGTTTCTCTTGCCTTTTT | TCTTCTGATTCTGCTGTGACAA |
| GULP1 | GCCCAAAGGAACAGAAGTTG | GGAATTTTCTGGCCTTCAGA |
| SLC401A | GCCAACTACCTGACCTCTGC | AAAGTGCCACATCCGATCTC |
| MSTN | TGACAGCAGTGATGGCTCTT | TTGGGTTTTCCTTCCACTTG |
| C2orf88 | CCTGTCCCAGGAAATCTTGA | GATGTTATTGCCTGCCCACT |
| STAT1 | TCTGCAGCTGTCTGAAGGAG | TGAATATTCCCCGACTGAGC |
| GLS | CCTAGCTTGGAAGATTTGCTG | CAGACGTTCGCAATCCTGTA |
| ZNF804A | ATGACCATGCTCACAAGCAG | TTTCGAGCAAATTCCCTTTG |
| COL5A2 | AGGAGAGAGAGGCCCAAAAG | CTCCATCAATTCCCTGAGGA |
| COL3A1 | CTGGTGCTAATGGTGCTCCT | TCTCCTTTGGCACCATTCTT |
| SOX11 | GTTCATGGTGTGGTCCAAGA | GCTGTCCTTCAGCATTTTCC |

**Table S1. List of primer sequences used in qPCR.**

| **Antibody** | **Company and Catalogue number** | **Dilution** |
| --- | --- | --- |
| Rabbit anti-SOX11 | Sigma; sab4200450 | 1:200 |
| Rabbit anti-COL3A1 | Abcam; ab7778 | 1:100 |
| Rabbit anti-RUNX2 | Santa-Cruz; sc10758 | 1:50 |
| Rabbit anti-STAT1 | Abcam; ab109320 | 1:200 |
| Goat anti-rabbit alexafluor 594 | ThermoFisher; A11012 | 1:200 |

**Table S2. Antibodies used in immunocytochemistry.**

| **Breed** | **Number of horses** | **Minor Allele Frequency (MAF)** | **Reference** |
| --- | --- | --- | --- |
| Arabian | 38 | 0.00 | Durward-Akhurst et al 2021 |
| Belgian | 20 | 0.00 | Durward-Akhurst et al 2021 |
| Clydesdale | 19 | 0.00 | Durward-Akhurst et al 2021 |
| Franches Montagne | 30 | 0.13 | Durward-Akhurst et al 2021 |
| Icelandic | 17 | 0.00 | Durward-Akhurst et al 2021 |
| Jeju pony | 21 | 0.02 | Durward-Akhurst et al 2021 |
| Morgan | 22 | 0.11 | Durward-Akhurst et al 2021 |
| Other (36 breeds) | 127 | 0.05 | Durward-Akhurst et al 2021 |
| Quarter horse | 103 | 0.66 | Durward-Akhurst et al 2021 |
| Shetland | 55 | 0.00 | Durward-Akhurst et al 2021 |
| Standardbred | 59 | 0.04 | Durward-Akhurst et al 2021 |
| Thoroughbred (US and European) | 76 | 0.53 | Durward-Akhurst et al 2021 |
| Welsh pony | 20 | 0.00 | Durward-Akhurst et al 2021 |
| >25 different breeds | 88 | 0.13 | Jagannathan et al 2019 |
| Thoroughbred (Japanese) | 101 | 0.50 | Tozaki et al 2021 |
| UK Thoroughbred fracture cases | 91 | 0.32 | N/A |
| UK Thoroughbred fracture controls | 86 | 0.22 | N/A |

**Table S3. Minor allele frequencies of the SNP in different breeds.**

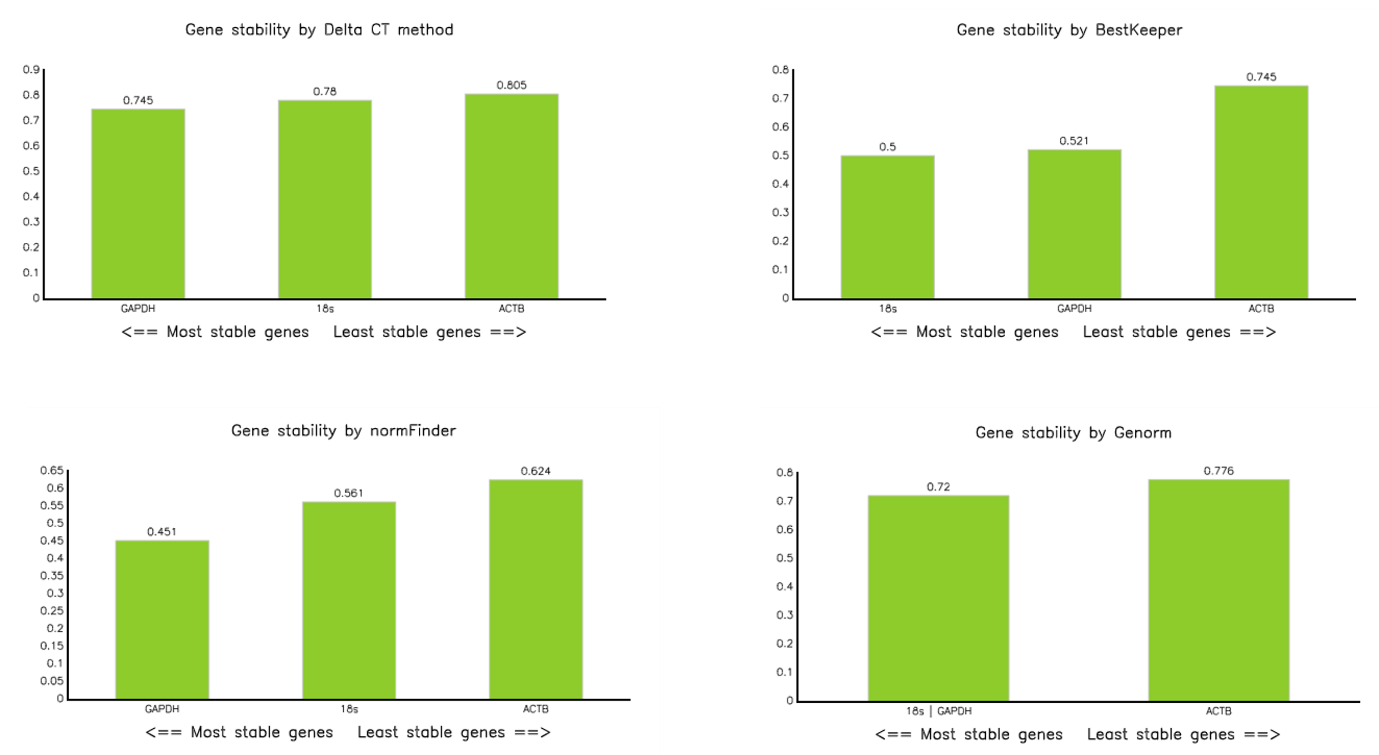

**Supplementary Figure 1.** Output from RefFinder showing the stability testing of three housekeeping genes demonstrated that 18S RNA and GAPDH were more stable than ACTB across four different housekeeping computational tools.

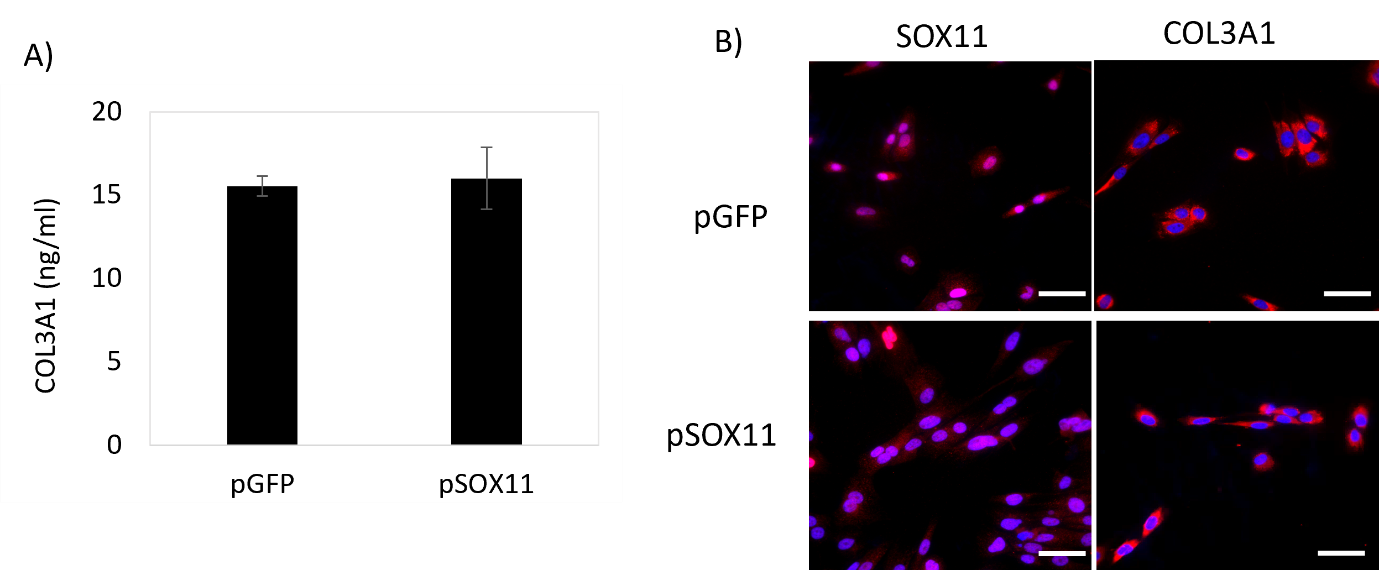

**Supplementary Figure 2.** COL3A1 protein levels do not change in response to SOX11 overexpression. A) No difference in total COL3A1 protein levels is detected using ELISA. Error bars represent the s.e.m from cells derived from three different donors. *p<0.05. B) SOX11 and COL3A1 (both shown in red) protein expression in control (pGFP) and SOX11 overexpressing (pSOX11) cells. DAPI staining of the nucleus is shown in blue. Scale bar = 40 µm. Images are representative of replicates using cells derived from three different donors.

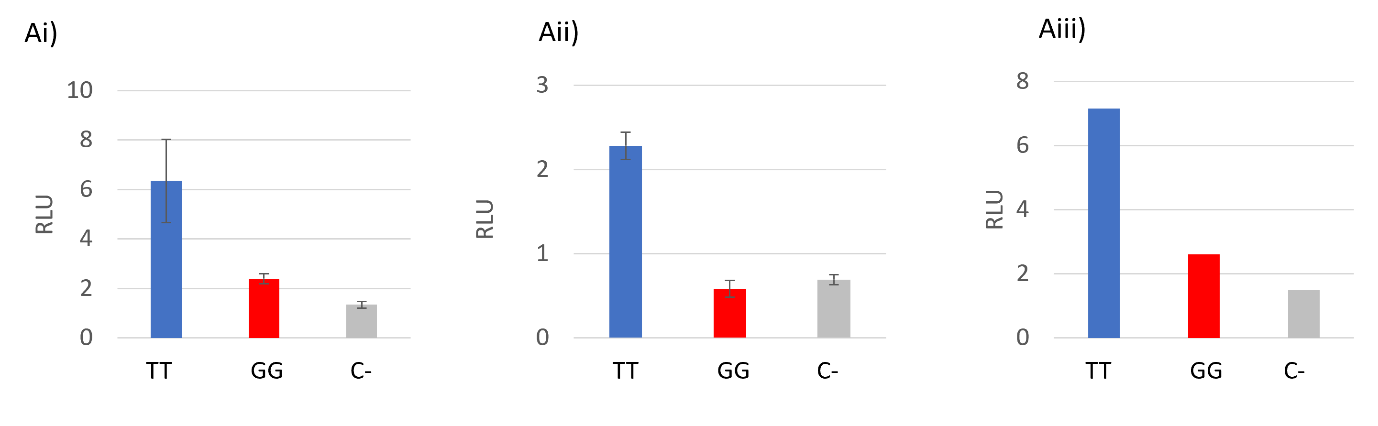

**Supplementary Figure 3.** Luciferase assays showing the relative light unit (RLU) for the 3220 bp region upstream of COL3A1 containing either the reference allele (TT) or the alternative allele (GG) compared to a promoterless control (C-). Ai – iii) show the results of independent luciferase assays set up on different days using skin cells derived from the same horse. Error bars represent the s.e.m of three to six technical replicates.
